# Landscape configuration of an Amazonian island-like ecosystem drives population structure and genetic diversity of a habitat-specialist bird

**DOI:** 10.1101/2020.12.25.424395

**Authors:** Camila D. Ritter, Camila C. Ribas, Juliana Menger, Sergio H. Borges, Christine D. Bacon, Jean P. Metzger, John Bates, Cintia Cornelius

## Abstract

**Context:** Amazonian white-sand ecosystems *(campinas)* are open vegetation patches which form a natural island-like system in a matrix of tropical rainforest. Due to their clear distinction from the surrounding matrix, the spatial characteristics of *campina* patches may affect the genetic diversity and composition of their specialized organisms such as the small and endemic passerine *Elaenia ruficeps.*

**Objectives:** Here, we estimate the relative contribution of the current extension, configuration and geographical context of *campina* patches to the patterns of genetic diversity and population structure of *E. ruficeps.*

**Methods:** We sampled individuals of *E. ruficeps* from three landscapes in Central Amazonia with contrasting *campina* spatial distribution, from landscapes with large and connected patches to landscapes with small and isolated patches. We estimate population structure, genetic diversity, and contemporary and historical migration within and among the three landscapes and used landscape metrics as predictor variables. Furthermore, we estimate genetic isolation by distance and resistance among individuals sampled within landscapes.

**Results:** We identified three genetically distinct populations with asymmetrical gene flow among landscapes and a decreasing migration rate with distance. Within each landscape, we found low genetic differentiation without genetic isolation by distance nor by resistance. In contrast, we found differentiation and spatial correlation between landscapes.

**Conclusions:** Our results uncover population dynamics of *E. ruficeps* through time. Together with previous studies, this suggests that both regional context and landscape structure shape the connectivity among populations of *campina* specialist birds, and that Amazonian landscapes, together with their associated biota, have responded to recent climatic changes.

## Introduction

Landscapes are mosaics of environments with distinct structure and biotic composition. Natural island-like systems such as habitat patches, caves, and mountaintops provide important contributions to landscape structure and diversity (Itescu 2019). Due to their well-defined borders and distinction from the surrounding habitats, the spatial characteristics of island-like systems may influence biological assemblages and their attributes including the genetic diversity and differentiation. These island-like systems can vary in the extent of insularity they impose on the taxa they harbor, affecting the extent to which organisms can disperse and colonize new patches (Itescu 2019).

Amazonia has the highest biodiversity among all tropical rainforests and is a global biodiversity hotspot (Hansen et al. 2013). The predominant view of Amazonia as a homogeneous humid tropical forest does not match the heterogeneity of landscapes it harbors (Myster 2016; Tuomisto et al. 2019). Indeed, Amazonia comprises diverse vegetation formations from humid tropical forests *(terra-firme)* to non-forested formations, such as white-sand grasslands and shrubby habitats (Anderson 1981; Adeney et al. 2016; Capurucho et al. 2020a).

White-sand shrub and grassland patches, hereafter *campinas*, are naturally fragmented and resemble islands in a “sea” of forests, growing on nutrient-poor soils (Prance 1996; Fine et al. 2010; Ritter et al. 2018; Costa et al. 2020; Capurucho et al. 2020a). *Campina* patches cover approximately 1.6% of the Amazon basin (Adeney et al. 2016), yet are an important Amazonian island-like system, harboring a unique biota (Borges et al. 2016a; Costa et al. 2020; Capurucho et al. 2020a). Landscapes with *campina* patches have different spatial configurations throughout Amazonia, composed of large and connected patches in the north and small and isolated patches in the south (Borges et al. 2016a).

Moreover, properties of *campina* landscapes, such as habitat amount, patch isolation and matrix properties, vary across space and time. As such, it is expected that gene flow among populations and hence genetic diversity of populations inhabiting *campina* patches will depend on the structure of these landscapes. Thus, landscapes with more *campina* habitat cover and with connected patches should harbor a higher genetic diversity than landscapes with reduced habitat and isolated patches. However, the effects of landscape configuration on the organisms that thrive in naturally fragmented *campinas* remain poorly understood (but see Capurucho et al. 2013; Borges et al. 2016a).

Several factors may restrict the movement of individuals in island-like systems, such as *campinas.*In naturally heterogeneous landscapes, restrictions of movement and gene flow can be due, for instance, to geographic distance (isolation by distance; Wright 1943), or to non-suitable habitat (isolation by resistance; e.g. McRae 2006; DiLeo and Wagner 2016). Dispersal promotes gene flow and connects geographically isolated populations, increases genetic diversity, and reduces inbreeding (Ronce 2007). However, dispersal through non-suitable habitats also represents high energetic costs and mortality risks (Fahrig 1998; Gruber and Henle 2008).

The geographical distance between patches, within and among landscapes, and the type and configuration of environments in the matrix may affect the ability of a species to disperse (Bates 2002). Different matrices create variable resistance to individuals’ movement (Itescu 2019). In Amazonia, white water rivers, such as the Amazon River, appear to impose large resistance for white-sand vegetation specialist birds (Capurucho et al. 2013; Matos et al. 2016; Ritter et al. 2020). However, little is known about how composition and configuration of *campina* landscapes shape movements of specialist species. Also, landscapes are dynamic over time and thus movement patterns of individuals may differ within and among landscapes over time (Manicacci et al. 1992). Understanding these movements through time provides information on landscape configuration changes in the past, contributing to the long debate about stability of Amazonian habitats during the Quaternary (Cheng et al. 2013; Wang et al. 2017; Bicudo et al. 2019; Rocha and Kaefer 2019).

Methods of molecular analyses have been successfully used to investigate patterns and to infer processes related to the origin and maintenance of biodiversity (e.g. Antonelli et al. 2018; Silva et al. 2019). The use of gene sequencing can reveal historical patterns through phylogeographic studies (Avise 2009). On the other hand, the genotyping of microsatellite markers can reveal contemporary patterns, because they are highly polymorphic due to their high mutational rate (Tautz 1989), and therefore have been considered ideal for studies of contemporary population structure (Frankham et al. 2002). In this context, the use of molecular markers with distinct evolutionary rates may uncover how the interaction between landscape features and micro-evolutionary processes shapes patterns of genetic structure and diversity in time and space (Capurucho et al. 2013).

In this study we investigate the effects of landscape configuration on population genetic structure and diversity in a white-sand vegetation specialist bird species restricted to Amazonian *campina* patches, *Elaenia ruficeps* (Aves: Tyrannidae; Rheindt, Norman and Christidis 2008; Borges et al. 2016b), employing mitochondrial gene sequences and microsatellite markers.

We address the following questions:

1. How do genetic diversity, population structure, and migration rates differ within and among three *campina* landscapes with contrasting configuration? We expect differences between genetic metrics measured through markers with faster (microsatellites) and slower (DNA sequence) evolutionary rates that responded to processes at different time scales, with microsatellites more congruent with current landscape structure.
2. How does habitat amount and isolation of patches within and among landscapes affect population genetic diversity in *E. ruficeps?* We expect that both metrics will be important but habitat amount will be the strongest factor explaining genetic diversity.
3. What is the relative importance of geographical distance and matrix resistance in limiting gene flow in *E. ruficeps?* We expected that habitat matrix resistance will better explain genetic differentiation among populations than geographic distance. We explicitly tested if *terrafirme forest* and rivers limited movement of *E. ruficeps* individuals more than other landscape matrix types, such as seasonally flooded forests.

## Materials and Methods

### Study area

We sampled birds in three landscapes (each *ca* 50 x 50 km) north of the Amazon River (Fig. 1A): Aracá (0°28’7.76” N, 63°28’32.20” W; Fig. 1B), Viruá (1°36’ N, 61°13’ W; Fig. 1C), and Uatumã (2°17’9.19” S, 58°51’53.92” W; Fig. 1D). The Aracá landscape lies on the eastern side of the middle part of the Negro river basin on the western margin of the Branco river (i.e. in the Branco-Negro interfluve) and has the highest *campina* vegetation coverage (45.33% of its area in 50 x 50 km^2^) distributed as large and connected patches. The Viruá landscape is located on the eastern margin of the Branco River, and has intermediate *campina* vegetation coverage (28.2 % of its area) distributed as both large and small interconnected patches. This is the only site with anthropogenic disturbance due to an interstate road and a few secondary non-paved roads among the sampling sites. The third and southernmost landscape, Uatumã, is located on the banks of the Uatumã river, inside the limits of the Uatumã Sustainable Development Reserve. The Uatumã landscape has less *campinas* coverage (0.8% of its area) with small and isolated *campina* patches (Fig. 1 B-D). We established six sampling sites within the Aracá landscape, six sampling sites within the Uatumã landscapes, and four sampling sites within the Viruá landscape, with a total of 16 sampling sites. These sites were distributed across landscapes in *campinas* vegetation with distances ranging from 5 to 44 km among them within landscapes, as described in Capurucho et al. (2013) and Borges et al. (2016a).

**Figure 1A) -.**
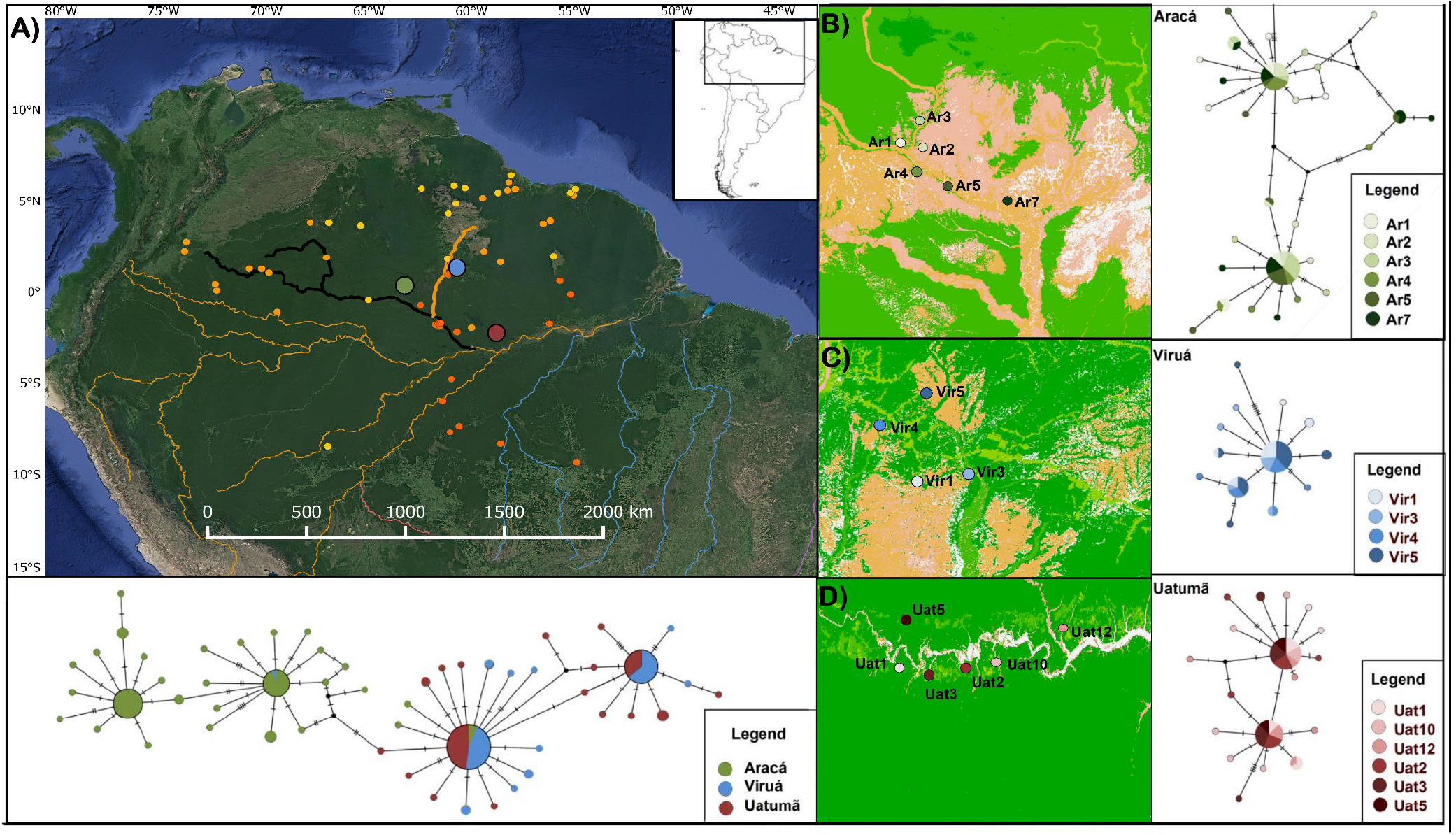
Map of the distribution of *Elaenia ruficeps.* Points in yellow are from the Global Biodiversity Information Facility (GBIF) public database (general and potentially biased by mis-identification); points in bright orange are from museum collections (highly curated locality information). Points in dark orange are areas with available tissue samples. Points in green, blue, and red are the landscapes sampled in this study (Aracá, Viruá, and Uatumã respectively). The main rivers of the Amazon basin are shown according with their water color; rivers with high sediment concentration are brown, with low sediment concentration are blue. We highlight the Negro river in Black and the Branco river in brown both with tick lines. B) shows in detail the sites sampled in Aracá with the respective haplotype network; C) shows the sites sampled in Viruá with the respective haplotype network and; and D) shows the sites sampled on Uatumã and the respective haplotype network.

### Landscape metrics

We used categorical maps with six pre-defined classes: *terra-firme forest, campinas, campinarana* (white-sand patches with taller vegetation coverage than *campinas*), flooded forest, water, and anthropogenic areas (see Capurucho et al. 2013 for details on the classification method). We used ArcGIS v.9.1 (Press 2005) and Fragstas v.3.4 (McGarigal et al. 2002) to calculate two landscapes metrics. The first was a habitat amount metric calculated as the area of *campina* vegetation in a radius of 5 km around each sampling site. As a configuration metric, we used the proximity index, an isolation measure of each patch in which sampling sites were located, that was based on the sum of the area of neighboring patches within a 5 km search radius, weighted by the distance to neighboring patches (Gustafson and Parker 1994). The 5 km radius was selected based on dispersal kernels described for several Neotropical bird species; most Amazonian birds disperse less than 5 km (Van Houtan et al. 2007). Habitat amount and proximity index were not correlated (Pearson correlation = 0.3, p = 0.24).

### Sampling

We defined sampling area as a 500 m radius circle centered at each sampling site, in which 20 mist-nets (12 m long, 36 mm mesh size) were equally distributed into four mist-net lines. In order to reduce the probability of sampling only one family group, at least two mist-net lines were moved each day to cover other parts of the sampling sites. Sampling was conducted during the dry season of 2010 and 2011, and each site was sampled as many days as needed to capture at least 10 individuals per site, ranging from 2-5 days per site, but in four sites the intended number of individuals was not attained (see Table S1). A blood sample (~ 50 μl) was taken from each captured individual, stored in ethanol, and deposited in the Genetic Resources Collection of the National Institute for Amazonian Research (INPA, Manaus, Brazil). Voucher specimens (maximum of five per landscape) were also collected and deposited at INPA Bird Collection (Table S1).

### DNA sequencing

DNA was extracted from blood or tissue samples using Promega DNA Purification Kit (A1125). The complete sequence of the mitochondrial NADH Dehydrogenase 2 (ND2) gene was amplified using the external primers L5204 and H6313 (Sorenson et al. 1999). For this study, we also designed a primer for reverse gene sequencing (H1031; 5’-TAGGATTGTAGGGGATAAAGGTA-3 ‘), because some samples did not amplify well with H6313. Amplification and sequencing details are described in Capurucho et al. (2013). Contiguous sequences were assembled and aligned in Geneious v. 5.6.5 (Biomatters 2012).

### Microsatellite genotyping

All individuals of *E. ruficeps* were genotyped at 15 microsatellite loci described in Ritter et al. (2014), using protocols and PCR conditions therein. PCR products were run on an ABI PRISM 3730 DNA Analyzer; size scoring was performed with GeneMarker^®^ v2.2.0 (Hulce et al. 2011). We calculated the number of alleles per locus, observed and expected heterozygosity (Ho and He), deviations from Hardy-Weinberg equilibrium (HWE), and the exact tests of linkage disequilibrium between pairs of loci for the three landscapes using Genepop Web v.4.2 (Raymond and Rousset 1995; Rousset 2008; Table S2).

### Genetic diversity, population structure, and migration rates

To investigate if genetic diversity varies within and among landscapes we calculated four genetic diversity metrics (two based on mitochondrial and two on microsatellite data). For each locality (both landscapes and sites within each landscape) we estimated the individuals nucleotide diversity (Pi) and haplotype diversity (HD) based on ND2 mitochondrial sequences using DnaSP v.5.10.01 (Librado and Rozas 2009). For the microsatellite data we estimated allelic richness using the rarefaction method implemented in the *Hierfstat* v.0.4.22 package (Goudet and Jombart 2015) in R v.3.2.5 (R Core Team 2015), and calculated the microsatellite genetic diversity (Theta) using Arlequin v.3.11 (Excoffier et al. 2005).

To describe historical population structure within and among landscapes, we constructed haplotype networks with ND2 sequences, with all individuals together and for individuals from each landscape separately, using a minimum spanning network (Clement et al. 2002) with Popart v.1.7 (Leigh and Bryant 2015). We used BAPS v.6.0 (Bayesian Analysis of Population Structure; (Corander et al. 2013) to infer the number of clusters *(K)* based on the mitochondrial data using all individuals. Likelihood values of the mixture analysis were calculated three times for each number *K* of subpopulations, ranging from 1 to 20 (since the sites number was 16 and we expected no more than 20 population), accepting the partition with *K* value with higher likelihood, which were run until achieving convergence.

To describe current population structure within and among landscapes, we used microsatellite data. We used Structure v.2.3.4 (Pritchard et al. 2000), to infer the number of genetically distinct populations (*K*). We assumed an admixture model with correlated allele frequencies and the LOCPRIOR model (Hubisz et al. 2009). We used two LOCPRIOR options, first, we made analyses at the landscape level, where Aracá, Viruá and Uatumã were considered each a single locality. Secondly, we analyzed the data using each sampling site as unique localities. To identify the best estimate of *K* from 1 to 20 (both sampling site as populations and landscapes as populations), we set a burn-in period of 100,000 followed by additional 1,000,000 iterations, and 20 replicates were run at each *K*. We determined *K* based on the log posterior probability of the data for a given *K*(Pritchard et al. 2000), and on the rate of change in the log probability of the data between successive clusters—the ΔK statistic (Evanno et al. 2005). These analyses were performed in Structure Harvester v.0.6.94 (Earl 2012). All runs were averaged at the best *K* with Clumpp v.1.1.2 (Jakobsson and Rosenberg 2007) and results were visualized with Distruct v.1.1 (Rosenberg 2004).

We inferred historical migration rates using mitochondrial sequences in Migrate-N v.3.6 (Beerli 2009). Under a coalescent framework and the infinite allele model, Migrate-N estimates migration rates (measured as a mutation-scaled immigration rate, M) up to ~4 effective population size (Ne) generations (thousands of years). We used slice sampling to run four statistically heated parallel chains (heated at 1.0, 1.5, 3.0, and 1,000,000) for 1,000,000 iterations, and excluded 100,000 iterations as burn-in. MCMC estimates of M were modeled with prior boundaries of 0 and 100,000. We used a full migration model and considered parameter estimates accurate when an effective sample size (ESS) >1,000 was observed (Converse et al. 2015). We multiplied the M by the mutation rate, 0.0105*10^-4 for the mitochondrial data (Lovette 2004; Weir and Schluter 2008). To test for spatial auto-correlation of migration rate, we performed a Mantel test with pairwise migration rates and geographic distances (Euclidean) using the *vegan* v. 2.4-3 (Oksanen et al. 2010) R package. We performed these analyses both between landscapes and between sampling sites separately.

To estimate current migration rates, we used the microsatellite data in BayesAss v.3.0 (Wilson and Rannala 2003), which applies a Bayesian approach and MCMC sampling to estimate migration (m) over the last few generations. This analysis was run with 10 million iterations, a sampling frequency of 2,000, a burn-in of 10%, and default settings. We estimated the migration rate between sampling sites and between the three landscapes separately.

Furthermore, to identify if past demographic changes can explain genetic diversity and migration rates, we inferred historical population demography using the Bayesian coalescent skyline plot method (Drummond et al. 2005) as implemented in Beast v.1.8.2 (Drummond et al. 2012). We chose the most suitable substitution model for the mitochondrial data based on Bayesian information criterion (BIC) with jModelTest2 v.2.1.10 (Darriba et al. 2012). We set the substitution model chosen by jModelTest2 (HKY + invariable sites) under a strict-clock model and the general avian substitution rate of mitochondrial evolution of 2.1% sequence divergence per million years (Lovette 2004; Weir and Schluter 2008). Runs of 100 million steps were performed, sampling every 10,000 steps under default settings. Skyline plots were constructed using Tracer v.1.6 (Rambaut and Drummond 2007). We reconstructed historical population size considering all populations together and then separately for Aracá from Viruá+ Uatumã following the populations identified with BAPS.

### Landscape metrics and genetic diversity

To investigate if landscape metrics predict genetic diversity metrics we calculated genetic metrics through nucleotide diversity (Pi) and haplotype diversity (HD) from mitochondrial sequences and allelic richness (NG) and genetic diversity (Theta) from microsatellite data. We calculated these metrics within each site and analyzed as a function of the two landscape metrics (habitat amount and proximity index) and of the landscape of origin of each site (Aracá, Viruá our Uatumã).

Thus for each dependent variable (Pi, HD, Theta, and NG), we defined a set of models to explain variation in genetic diversity. The final model set included models for each single landscape metric, and additional models with additive and interaction terms of the landscape origin to determine whether landscape context was also an important factor (i.e. to which landscape each group of sampling sites belongs to). The final model set also included a constant, intercept-only model, comprising a total of seven models for each dependent variable (Table S3).

Models were selected using an information theory approach based on AIC (Akaike 1974) and using the corrected AIC (AICc) for small sample sizes (Burnham and Anderson 2002). Models with dAIC =< 2 were considered equally plausible and we used the normalized model weight (wi) to contrast the best model to the constant (no-effect) model. We used generalized linear models (Crawley 2007) with Gaussian error distribution after checking for the distribution of residuals. Before run the analysis landscape metrics were standard to mean = 0 and variance = 1 to make different metrics comparable. The GLM analyses were performed using the *vegan* v. 2.4-3 (Oksanen et al. 2010) package and the model selection was made using the *bbmle* v.1.0.20 (Bolker and Bolker 2017) package, both in R.

### Geographical isolation by distance and by resistance

To determine if genetic differentiation is better predicted by geographical distance or resistance we calculated the pairwise genetic differentiation F_ST_ (Weir and Cockerham 1984) for both ND2 and microsatellite data between landscapes and among sites within landscapes separately using the *fstat* function in the *Hierfstat* v.0.5.7 R package, with 1,000 permutations to obtain significance (Goudet 2001). To investigate patterns of isolation by geographical distance, we performed Mantel tests in the *vegan* v. 2.4-3 R package. We used a pairwise geographic (Euclidean) and a pairwise genetic distance (F_ST_ values). We performed these analyses both between landscapes and between sampling sites within each landscape separately.

To investigate the patterns of isolation by resistance we assigned resistance values to vegetation cover within each landscape based on a questionnaire given to four expert Amazonian ornithologists for each landscape category for *E. ruficeps.* Values ranged from 0.01 (less resistance) to 0.99 (more resistance). We took the average resistance value of each landscape category to calculate the isolation by resistance (Table S3). We used the *gdistance* v. 1.2-2 (Etten 2017) R package to create the transition layer using the inverse of the sum of each pixel to create the conductance layer (Fig. S1) and the *commuteDistance* function that calculates the expected random-walk commute resistance between nodes in a graph, to create the pairwise resistance matrix for each landscape. We then performed a Mantel test using the pairwise genetic distance (F_ST_ values) against the resistance distance. Additionally, we calculated the minimum resistance distance (i.e, least cost path) for each pair of sites and a Mantel test with the pairwise genetic distances (F_ST_ values).

## Results

### Genetic diversity, structure, and migration

We obtained 978 bp of the ND2 gene, yielding a total of 178 sequences with 62 variable sites. Haplotype diversity from mitochondrial data of all samples was 0.79 (+/− 0.08) and nucleotide diversity was 0.002 (+/− 0.001). For microsatellite data, we scored the same 178 individuals at 15 loci. No departure of Hardy-Weinberg Equilibrium was detected at any locus and no pair of loci was in linkage disequilibrium (see Table S2 for number of alleles per locus). Aracá landscape had the highest haplotype (0.84 +/− 0.07) and nucleotide diversity (0.003 +/− 0.0006) for mitochondrial data. For microsatellite data Aracá also had the highest allelic richness (24 +/− 3.58) but Viruá had the highest genetic diversity (1.69 +/− 0.04, Table S4).

We detected low but significant genetic differentiation among landscapes for both mitochondrial and microsatellite data (Table 1). For mitochondrial data, Viruá and Aracá had the largest differentiation (F_ST_ = 0.1, p < 0.05), while the largest differentiation for microsatellite data was inferred between Viruá and the Uatumã landscapes (F_ST_= 0.2, p < 0.05, Table 1). Comparing among all sampling sites, within and among landscapes, mitochondrial results revealed low but significant differentiation among almost all sites within each landscape, except among some Aracá sites (Table S5). Genetic differentiation among sites from different landscapes was highest than sites within landscapes and significant, except between some Uatumã and Viruá sites (Table S5). Microsatellite results revealed several cases of non-significant differentiation among sites, within and among landscapes (Table S6). For sites in different landscapes, the sites from Uatumã were more differentiated than sites of both Aracá and Viruá (Table S6).

**Table 1 -.**
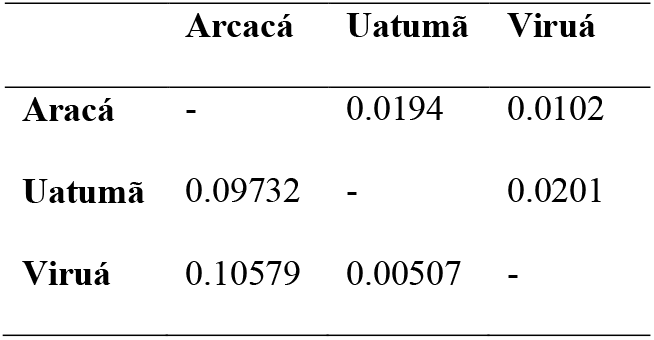
F_ST_ among landscapes. Values above the diagonal are microsatellite F_ST_ and below the diagonal are ND2 sequence F_ST_. All values are significant at P < 0.05.

We found 56 mitochondrial haplotypes that grouped into three main clusters. Most haplotypes from Aracá landscape were not shared with the Viruá and Uatumã landscapes, and within Aracá the haplotypes were grouped in two main clusters. Only one Aracá haplotype was shared with the other two landscapes (Fig. 1A). Despite clear differentiation between Aracá and the other two landscapes, the haplotype networks inside each landscape had little or no small-scale geographic structuring. Within each landscape, sampled haplotypes occurred in almost all sampled sites (Fig. 1B-D). BAPS results agree with the haplotype networks and inferred *K*= 3 populations, with two groups within Aracá and one with all haplotypes from Viruá and Uatumã, including three Aracá haplotypes found in five individuals (Fig. 2A), log (marginal likelihood) of optimal partition = - 920.3443, 100% probability of K = 3. For microsatellites, the highest log posterior probability of the data and the highest value for *ΔK* obtained via Structure analysis also inferred *K*= 3 (Fig. 2B), however the populations recovered by the microsatellite data corresponded to the three sampled landscapes.

**Figure 2 -.**
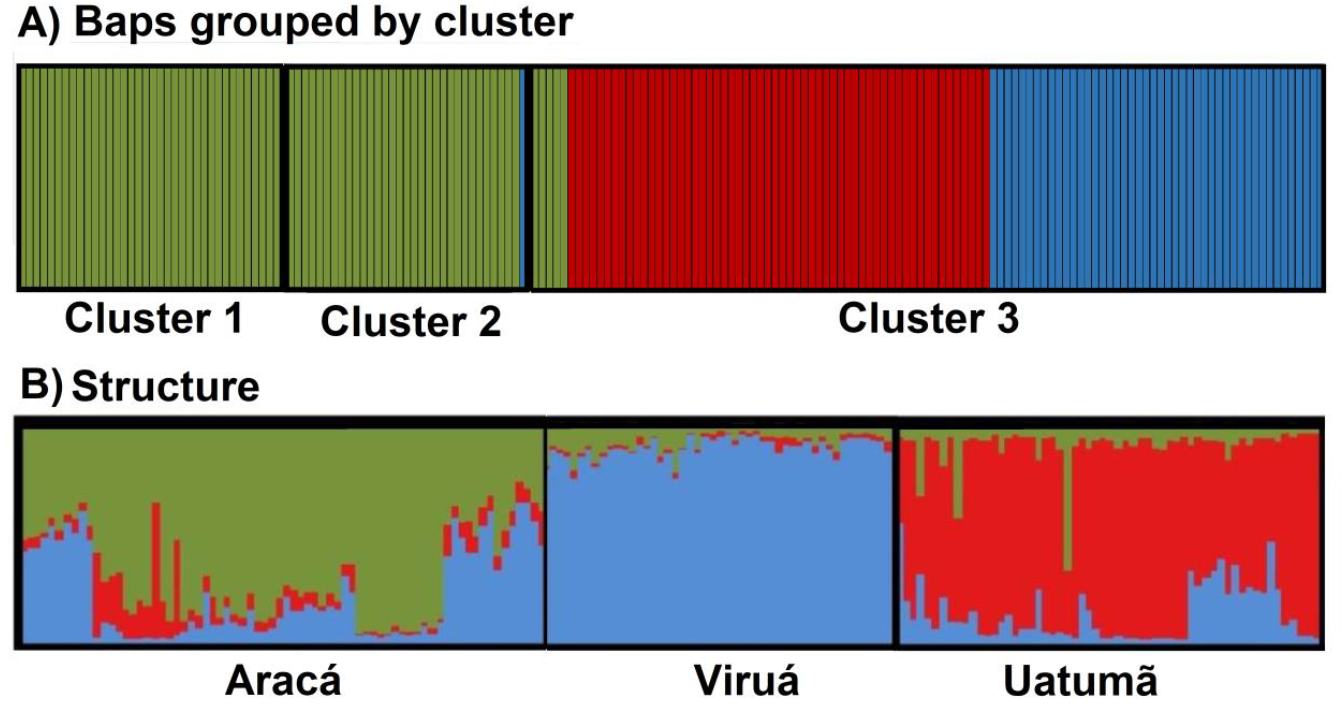
Population structure of *Elaenia ruficeps* based on (A) mtDNA (BAPS) with individuals ordered by cluster membership and colored by landscape of origin (landscapes in decreasing order of *campina* habitat coverage: Aracá = green, Viruá= blue, Uatumã= red), and (B) microsatelites (Structure), for which clusters match the different landscapes. In both analyses recorded *K* = 3 genetic clusters, which are delimited by thick black lines.

Estimates of historical migration obtained from Migrate-N with mitochondrial data indicated low and asymmetrical gene flow from Uatumã to Viruá (0.0009) and from Viruá to Uatumã (0.0003), with even lower but symmetrical rates between Viruá and Aracá (0.0001 in both directions), and very low rates between Uatumã and Aracá (<0.00006 in both directions; Fig 3A). Estimates of contemporary migration obtained from BayesAss with microsatellite data resulted in high self-recruitment rates for all three landscapes (Aracá= 0.99 [±0.006], Uatumã= 0.67 [±0.005] and Viruá= 0.67 [±0.006]). Contemporary migration was also asymmetrical, with individuals moving mainly from Uatumã and Viruá towards Aracá, 0.32 (±0.008) and 0.32 (±0.009), respectively (Fig. 3B). Among all sites, historical (r = 0, p = 0.5) and contemporary (r = −0.08, p = 0.84) migration rates were not related with geographic distance (Fig. S2A). Also, among landscapes, contemporary (r = −0.03, p = 0.67) and historical (r = 0.16, p = 0.67) migration rates were not significantly related to geographical distance (Fig S2B).

**Figure 3 -.**
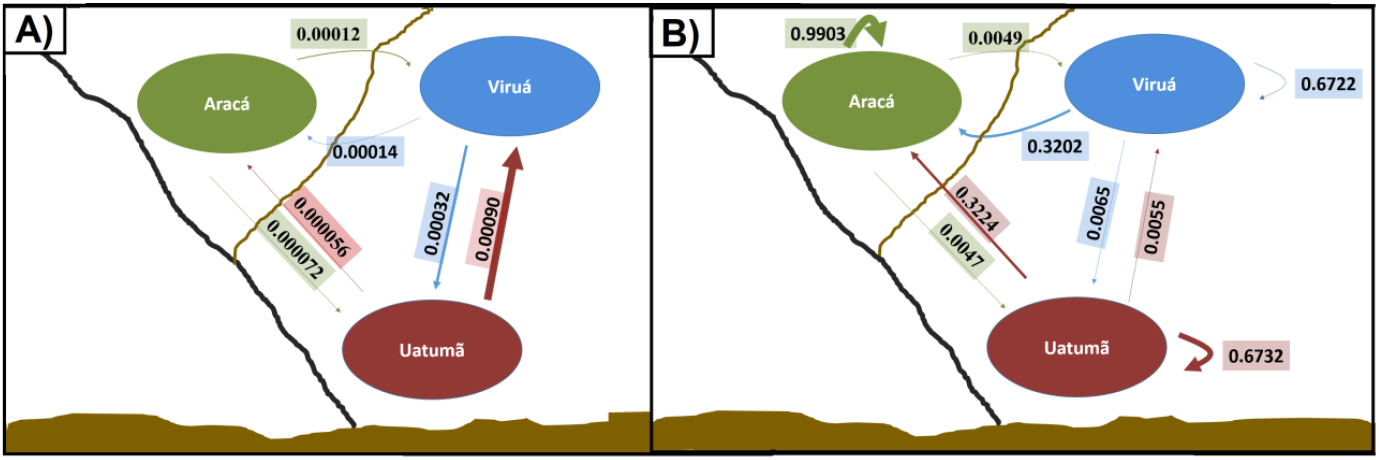
Pairwise migration rates. A) Historical migration rate calculated for mtDNA ND2 sequences in Migrate-N. B) Contemporary migration rate calculated in BayesAss using microsatellite data (CM microsatellite). The size of the arrows is proportional to the migration estimates. Black line represents Negro River and brow line the Branco River. Brown coloration in the bottom of figure represents the Amazon River. Historical migration shows the highest migration rate between Uatumã and Viruá, while contemporary migration shows higher migration from Uatumã and Viruá to Aracá.

Finally, analyses based on the mitochondrial data showed demographic expansion for *E. ruficeps* population as a whole. Bayesian skyline plot estimates showed general population expansion over the last 50,000 years (Fig. S3A). When we estimated demography separately, the Aracá population showed the demographic expansion over the last 50,000 years (Fig. S3B), but Viruá and Uatumã populations have maintained their population size constant over time (Fig. S3C).

### Landscape metrics and genetic diversity

For the nucleotide diversity metric (Pi), a single best model was selected that contained landscape of origin as the single predictor variable (AIC_w_ = 0.7686), while for haplotype diversity (H_D_) the single best model was the constant intercept-only model (AIC_w_ = 0.5573). For microsatellite genetic diversity (Theta), a single best model was selected that contained just landscape of origin as predictor variable (AIC_w_ = 0.732) and for allelic richness (NG) the single best model contained de proximity index as the single best predictor variable (AIC_w_ = 0.9376, Fig. 4A-D). See respective dAICc and AIC weights in Table 2 and best models estimated parameters in Table S7.

**Figure 4 -.**
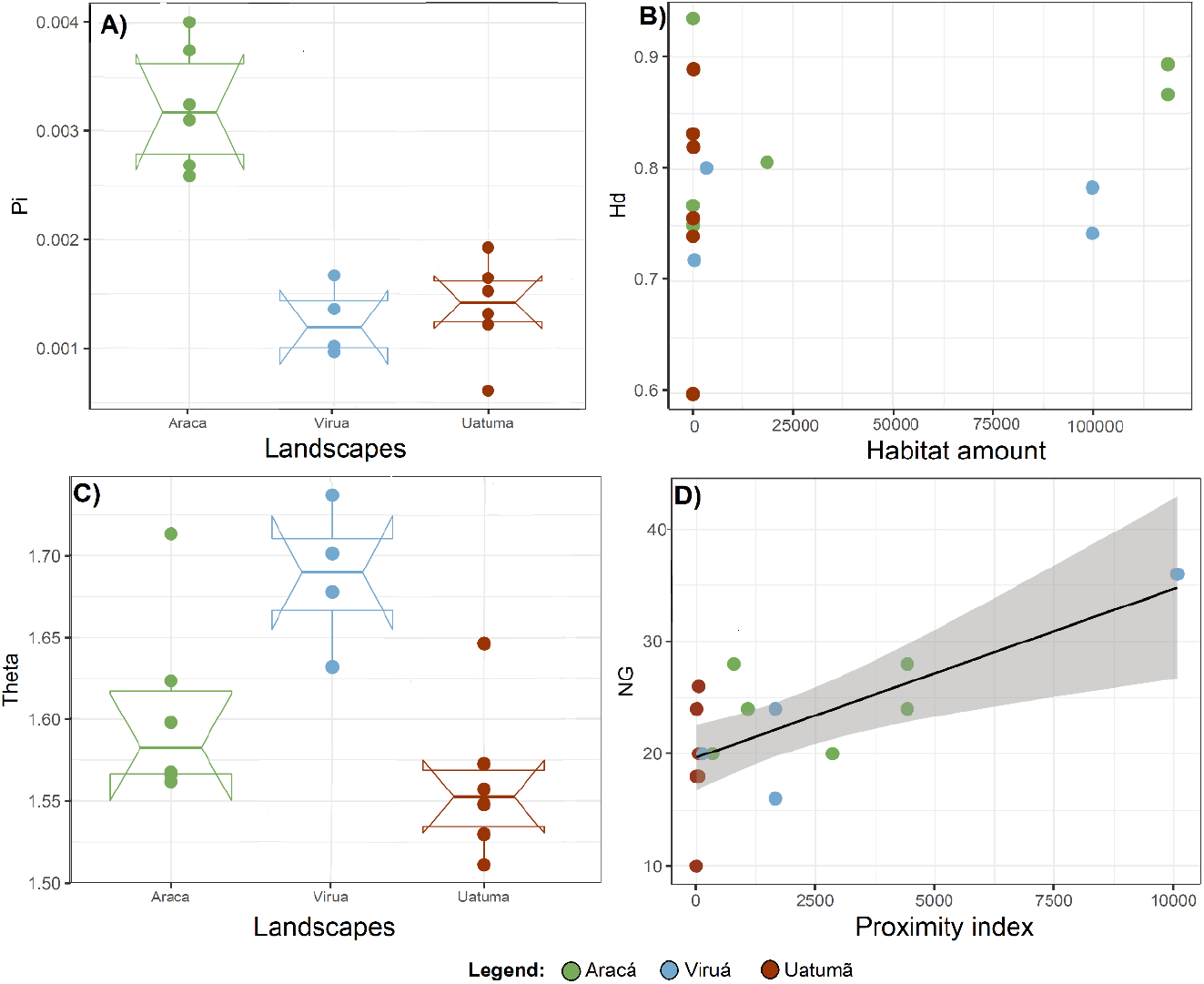
Best models about the source of variation of mitochondrial (A and B) and microsatellite (C and D) genetic diversity among sites within landscapes (Aracá = green, Viruá= blue, Uatumã = red). A) nucleotide diversity (Pi) based on ND2 is best explained by landscape; B) haplotype diversity based on ND2 sequences and habitat amount in m^2^ (but none of the predictor variables explained haplotype diversity; the constant model was selected as the best model). C) Theta from microsatellite data is best explained by landscape, and D) the microsatellite genetic diversity (NG) is best explained by the Proximity index.

**Table 2 -.** Variables used in model selection with their respective delta dAICc and weight values. The best fit model has a dAICc = 0 is present in bold as the alternative good models (dAICc =< 2). The genetic diversity variables for mitochondrial data are nucleotide diversity (Pi) and haplotype diversity and for the microsatellite data are Theta and NG. The independent variables are Habitat amount and Proximity index. The model used landscape as a fixed factor or as interacting variable.

| Model | Variables | Pi |  | H<sub>D</sub> |  | Theta |  | NG |  |
| --- | --- | --- | --- | --- | --- | --- | --- | --- | --- |
|  |  | dAICc | Weight | dAICc | Weight | dAICc | Weight | dAICc | Weight |
| M0 | Constant | 21 | <0.001 | **0** | **0.5573** | 5.4 | 0.049 | 6.9 | 0.0294 |
| M1 | Landscape | **0** | **0.7686** | 3.6 | 0.093 | **0** | **0.732** | 10.9 | 0.0039 |
| M2 | Habitat amount | 23.7 | <0.001 | 2.1 | 0.1981 | 7.7 | 0.016 | 9.8 | 0.0069 |
| M3 | Proximity | 23.3 | <0.001 | 3 | 0.1226 | 6.7 | 0.025 | **0** | **0.9376** |
| M4 | Landscape + habitat amount | 4.1 | 0.0986 | 7.1 | 0.0157 | 4.1 | 0.093 | 15.2 | <0.001 |
| M5 | Landscape * habitat amount | 13.2 | 0.001 | 12.5 | 0.0011 | 15.6 | <0.001 | 24 | <0.001 |
| M6 | Landscape + Proximity | 3.5 | 0.1307 | 7.7 | 0.012 | 4.3 | 0.084 | 7.6 | 0.0213 |
| M7 | Landscape * Proximity | 13.2 | 0.0011 | 16 | <0.001 | 13.7 | <0.001 | 15.9 | <0.001 |

### Geographical distance, resistance and gene flow

Genetic distance (F_ST_) for both microsatellite (r = 0.41, p = 0.01) and mitochondrial (r = 0.48, p = 0.001) data were positively correlated with geographic distance among landscapes (Fig 5A and B). However, no correlation with geographic distance was found among sites within each landscape (Table S8; Fig. 5C and D). No significant relationship was found between genetic differentiation (F_ST_) and resistance between sites within each landscape, in either dataset (mitochondrial or microsatellite) using the random-walk commute resistance (Table S8; Fig. 5E and F) and the pairwise minimal resistance between the sites.

**Figure 5 -.**
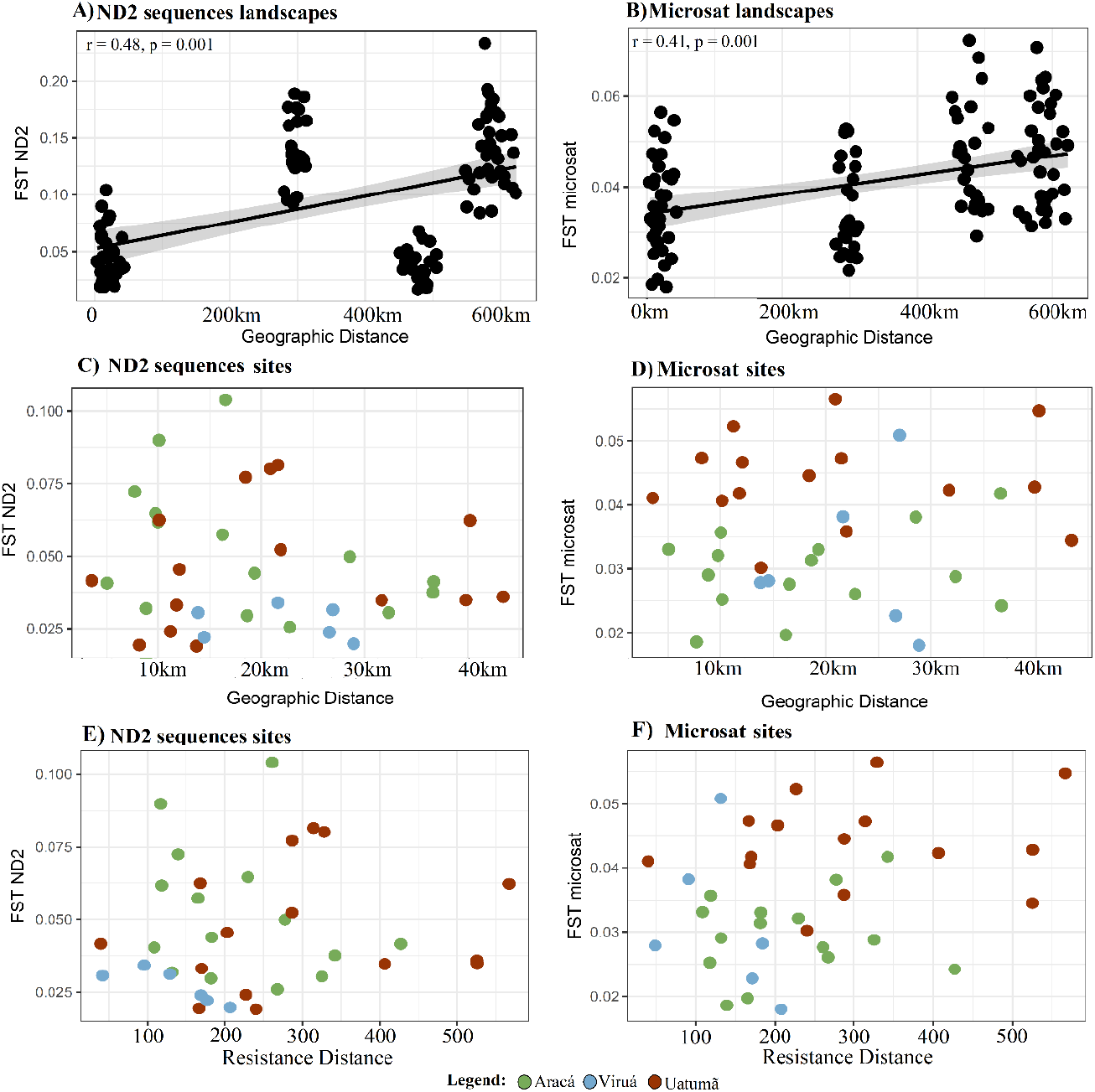
Pairwise genetic distance (F_ST_) and geographical distance relationship calculated with Mantel test. A) F_ST_ from ND2 sequence data and B) microsatellite data both plotted against the geographical distance between landscapes. C) F_ST_ from ND2 sequence data and D) microsatellite data both plotted against the geographical distance between sites inside each landscape. E) F_ST_ from ND2 sequence data and F) microsatellite data both plotted against the resistance distance between sites inside each landscape. The geographical distance is just significant between landscapes. Above pairwise matrix for all sites and below the data from each landscape: green = Aracá; blue = Viruá and; red = Uatumã.

## Discussion

We used a set of molecular markers with different evolutionary rates to analyze patterns of genetic diversity and population structure of *Elaenia ruficeps*, a white-sand specialist bird, by sampling three landscapes with different amount and configuration of *campina* patches in central Amazonia, we found that: 1) landscapes harbor genetically distinct populations, with asymmetrical gene flow among them; 2) historical and contemporary estimates of genetic structure and migration rates differ, implying dynamic connections among landscapes through time; 3) overall genetic structure (diversity and differentiation) is best explained by a regional effect (i.e. landscape of origin), than by habitat configuration, except for allelic richness which increases with patch proximity (more connectivity), supporting some evidence for local movement restriction between isolated patches; and 4) genetic differentiation increases with geographical distance among landscapes, whereas within landscapes no isolation by distance or by resistance is detected although small genetic differentiations is detected among patches. That suggests that dispersal of *E. ruficeps* between *campina* patches is restricted at least to some degree locally, but gene flow may be much more hampered by dispersal limitation at regional scales (between landscapes). Thus, our results stress the high complexity in *E. ruficeps* population dynamics in a habitat with insular nature.

### Genetic diversity and population structure: historical influences

The landscape with greatest habitat amount, Aracá, had the highest mitochondrial nucleotide (Pi) and haplotype (HD) diversity, and two mtDNA populations recovered in population structure analyses (Fig. 1A; Fig. 2A). A similar pattern of high genetic diversity and population structure was found for another white-sand specialist bird, *Xenopipo atronitens*, in the same region (Capurucho et al. 2013). Thus, it is likely that historical landscape alterations, such as glacial cycles, may have caused past population isolation within the Aracá landscape.

Population expansion starting around 50,000 years before present (Fig. S2) was found for *E. ruficeps* in our analyses, in agreement with other Amazonian bird species from *campinas*(Capurucho et al. 2013; Matos et al. 2016), but in contrast to results obtained for the same species using two additional nuclear markers (Ritter et al. 2020). This difference may be due to the lower mutation rates of nuclear markers (Allio et al. 2017), and increased sampling per locality used here. These historical demographic changes indicate that the populations of *E. ruficeps* may have started expanding in the last inter glacial, before the Last Glacial Maximum (LGM; Clark et al. 2009). When demography was estimated separately for each landscape, Aracá showed demographic expansion over the last 50,000 years (Fig. S3B), whereas Viruá+ Uatumã showed constant population size (Fig. S3C), suggesting that glacial cycles incurred variable impact in different regions of Amazonia and may explain the highest Pi, due to population expansion, in Aracá.

Studies on both northern (Carneiro Filho et al. 2002; Horbe, Horbe and Suguio 2004; Teeuw and Rhodes 2004; Zular et al. 2019) and southern (Latrubesse 2002) Amazonian *campinas* indicate that this habitat responded to historical changes in climate, with the strongest signal detected in the north. An increase in sediment deposition, primarily from the Tepuis, and aeolian activity, on northern *campinas* (Teeuw and Rhodes 2004; Zular et al. 2019), could have increased connectivity among populations of white-sand specialist species by increasing the area and connectivity of campinas, and consequently increasing population size and genetic diversity, during drier climatic periods in the Aracá region. Contrastingly, the Viruá landscape currently has the highest Theta diversity. *Campina* patches in Viruá also have higher diversity of white-sand specialist bird species, possibly due to its proximity to other open habitat types such as the Northern South America savannas (Fig. 1; Borges et al. 2016a; Capurucho et al. 2020a).

Estimated migration rates were asymmetrical, as found for other Amazonian birds (Capurucho et al. 2013; Menger et al. 2017), and we also found distinct values for historical and contemporary migration. Historical migration was higher between Uatumã and Viruá, with rates from Uatumã to Viruá three-fold higher than from Viruá to Uatumã. The Aracá landscape appears to be historically isolated from the other two landscapes. The historical isolation of Aracá may be explained by alterations to its overall size and/or connectivity during the Pleistocene glacial cycles (Teeuw and Rhodes 2004) and by the Branco River formation (Cremon et al. 2016). The Branco River is a white-water river that separates Aracá from Uatumã and Viruá and, together with its floodplains covered by seasonally flooded várzea vegetation, appear to have a stronger resistance for *campina’s* specialist birds (Capurucho et al. 2013; Matos et al. 2016). Furthermore, as suggested by haplotype network and migration rates, both previously (Ritter et al. 2020) and in this study, this river is also a barrier for *E. ruficeps*, which may have limited historical migration for Aracá populations. Analyses of sedimentary deposits and regional geomorphology suggested that a long segment of the Branco River was established in Late Pleistocene (Cremon et al. 2016). The initial establishment of the Branco River at about 30 kya may have collaborated to increased isolation of the Aracá population, but with gradual development of floodplain vegetation the barrier effect may be less pronounced since then. It is also possible that with the *terra-firme* canopy cover becoming less dense during past drier periods (Cowling et al. 2001), as hypothesized for northern Amazonia during the LGM (Häggi et al. 2017), the forested matrix surrounding *campinas* may have also turned more permeable than flooded forests along the Branco River, allowing for larger migration between Viruá and Uatumã, while Aracá remained isolated.

In summary, Pleistocene glacial cycles are a likely driver of population dynamics in of *E. ruficeps* through the increase of individual mobility across *terra-firme* forests in dry periods, while in the more isolated Aracá landscape, the continuous availability of the white-sand areas, even in wetter periods, may explain the higher genetic diversity. Genetic diversity patterns found for *E. ruficeps*are congruent with findings from other white-sand specialist birds (Capurucho et al. 2013; Matos et al. 2016), corroborating the idea that Pleistocene glacial cycles have deeply influenced Amazonian biogeographical history, and shaped current inter and intra-specific diversity (Rangel et al. 2018). This combined evidence from white-sand specialist birds suggests a dynamic interaction between closed canopy forests, open forests and non-forest / open vegetation areas (Cowling et al 2011; Arruda et al 2017), indicating that past climatic change deeply influenced Amazonian biogeographic history, and contradicting previous suggestions of a stable landscape in Amazonia during the Quaternary (Smith et al 2014).

### Genetic diversity and population structure: contemporary influences

In contrast to the historical scenario, microsatellite data indicate that current migration occurs primarily from Uatumã and Viruá towards Aracá, with lower migration rates in all other directions. Asymmetrical gene-flow arises due to more favorable dispersal conditions in one direction or due to source-sink dynamics across heterogeneous environments (e.g., Oswald et al. 2017; Hauser et al. 2019; Moussy et al. 2018). Aracá has the largest area of *campina* vegetation and is the most internally connected landscape. Furthermore, Aracá has in general the largest genetic diversity as measured here by three of the four indices, and in this context Aracá could function as a source population with a higher rate of emigration from Aracá towards the other populations. However, we found the opposite pattern, a higher migration rate towards Aracá, the largest and more connected population.

Considering the recent population expansion documented in Aracá over the last 50,000 years, in contrast to stability of population sizes in Uatumã and Viruá, it is possible that dispersal of individuals towards Aracá may be the result of emigration from small *campina* patches with little resource availability (e.g., Uatumã) or from landscapes that have been more affected by human impact (e.g., Viruá) with overall lower carrying capacity, but that are still able to maintain stable populations and thus are probably not sinks. Therefore, the asymmetrical gene flow in our study is most likely not consistent with a source-sink dynamic, and other mechanism should be investigated. An increased cost for dispersing towards one direction, as observed along elevational gradients (Cheviron and Brumfield 2009) is unlikely in our study system, but it is possible that environmental fluctuations are less strong in northern Amazonia (Jimenez and Takahashi 2019), leading to more constant resource supply in Aracá (the northern most landscape).

Landscape structure and landscape features have been shown to be important in shaping genetic diversity at the local scale for Amazonian vertebrates (e.g., Bates 2002; Capurucho et al. 2013; Menger et al. 2018; Silva et al. 2020). Here we show that allelic richness (NG) decreased in more isolated *campina* patches, but with no effect of habitat amount, in contrast to other findings showing that habitat amount best predicts genetic diversity and species diversity in white-sand specialist bird communities (Capurucho et al. 2013, Borges et al. 2016a).

This suggests that current local movements of *E. ruficeps* are, at least to certain degree, shaped by the configuration of *campina* patches. However, for *Xenopipo atronitens* another white-sand specialist bird, haplotype and nucleotide diversity increased with the amount of habitat available, with no effect of configuration (Capurucho et al. 2013). This difference may be explained by different species traits and habitat use patterns, since *X. atronitens* individuals also use *campinarana*, those white-sand patches that have taller vegetation than *campinas*, and eventually exploit black-water flooded forests as well (Oren 1981; Ridgely and Tudor 2009). In contrast, *E. ruficeps* is more restricted to *campina* vegetation (Borges et al. 2016b). Additionally, *E. ruficeps*has a lower handwing index than *X. atronitens*, a trait that was found to be correlated with overall range size in white-sand specialist birds (Capurucho et al. 2020b). These differences in habitat use highlight the importance of considering species traits when addressing congruence in biogeographical scenarios (Papadopoulou and Knowles 2016). Thus, we conclude that white-sand specialist birds are affected by landscape structure, but different components of these landscapes influence movement patterns of different species and both habitat amount (for *X. atronitens*; Capurucho et al 2013) and configuration (for *E. ruficeps;* this study) appear to be important for driving spatial patterns of genetic diversity of these white-sand specialist birds.

Genetic distance among landscapes increased with larger geographic distances in both mitochondrial and microsatellite data. Although significant genetic differentiation was found among most sampling sites within landscapes, no pattern of isolation by distance or resistance was observed. More refined studies on habitat permeability for white-sand vegetation birds are needed to develop more accurate isolation by resistance models. Our results suggest that although dispersal ability of *E. ruficeps* is at least to certain degree restricted through intervening vegetation types (Ritter et al. 2020), it is still greater than overall dispersal ability for most *terra-firme* forest birds (Menger et al. 2017; Menger et al. 2018), but dispersal ability of *E. ruficeps* is lower when compared to dispersal of savanna birds (Bates et al. 2003; Ritter et al. 2020). In a previous study comparing the population structure of *E. ruficeps* with its sister species *E. cristata*, it was evident that *E. cristata* populations that occur in savanna, have a low population structure indicating a higher mobility than *E. ruficeps* (Ritter et al. 2020). Furthermore, birds that occur in the *terra-firme* forest are generally limited by geographic distance (e.g. Menger et al. 2017; Menger et al. 2018), while typical Amazonian open area (savannas) bird species appear to have low population genetic structure, even at large geographic distances and across biogeographical barriers (Bates et al. 2003; Ritter et al. 2020).

## Conclusions

Here, we infer population structure, genetic diversity and migration within *E. ruficeps*, a white-sand specialist bird, among three Amazonian landscapes, using both, mitochondrial and microsatellite data. Distinct population structure was found for the different markers used, indicating differences in historical and current patterns of connectivity among landscapes. Migration rates were asymmetrical and also indicated a distinct scenario in the past compared to current rates. Patch isolation within and among landscapes was important to explain spatial patterns of microsatellite genetic diversity (NG). Geographical distance limited dispersal among but not within landscapes. These results suggest that both current landscape structure and the history of *campina* patches determine genetic diversity patterns of *campina* specialist birds. Further studies will certainly foster our understanding of how biotic communities associated to white-sand patches are influenced by current and historical processes in Amazonia, considering that these communities will probably be affected by future climatic changes.

## Acknowledgments

We thank the Brazilian authorities, ICMBio (20524-3 ICMBio/MMA), CEUC-AM, PARNA Viruá, and RDS Uatumã (24597-2 ICMBio/MMA) for providing the collecting permits and logistical support, and Fundação Vitória Amazônica (FVA), and Instituto de Conservação e Desenvolvimento Sustentável do Amazonas (Idesam) for logistical support. Laboratory work was conducted at the Laboratório Temático de Biologia Molecular (LTBM-INPA). Genetic data were gathered in the Pritzker Laboratory for Molecular Systematics and Evolution at the Field Museum of Natural History (FMNH). We thank Arielle Machado to read an initial version of the manuscript and Josué Azevedo for help with isolation by resistance analysis. We thank Gisiane Lima, João Capurucho, and Thiago Laranjeiras to answer the questionnaire about matrix resistance values.

## Declarations

### Ethical approvals

ICMBio, CEUC-AM (20524-3 ICMBio/MMA), PARNA Viruá, and RDS Uatumã (24597-2 ICMBio/MMA) for collecting permits.

### Consent to participate

All authors declare to consent to participate of this study.

### Consent for publication

All authors declare to consent the publication of this study.

### Availability of data material

GenBank accession ND2 sequences ID: 2304423. Microsatellite data is available in Ritter *et al.* 2014.

### Code availability

Not applicable.

### Authors contribution

C.C., C.C.R., C.D.R., and S.H.B. designed the study. C.D.R. and J.M. generated and analyzed the data. C.D.R. wrote the manuscript with contribution of C.C., C.C.R., C.B., J.M., J.P.M., J.B., and S.H.B.

