## Supplementary Material for "Landscape configuration of an Amazonian island-like ecosystem drives population structure and genetic diversity of a habitat-specialist bird"

**Supplemantery Material for:**

**Table S1** – Accession number of samples used in this study with the Landscape and site were each sample was collected. The geographical coordinates are also provided.

| **Accession** | **Landscape** | **Site** | **Latitude** | **Longitude** |
| --- | --- | --- | --- | --- |
| A3733 | Aracá | AR1 | 0.5556 | -63.4983 |
| A3735 | Aracá | AR1 | 0.5556 | -62.4983 |
| A3737 | Aracá | AR1 | 0.5556 | -61.4983 |
| A3738 | Aracá | AR1 | 0.5556 | -60.4983 |
| A3740 | Aracá | AR1 | 0.5556 | -59.4983 |
| A3742 | Aracá | AR1 | 0.5556 | -58.4983 |
| A3217 | Aracá | AR1 | 0.5556 | -57.4983 |
| A3219 | Aracá | AR1 | 0.5556 | -56.4983 |
| A3732 | Aracá | AR1 | 0.5556 | -55.4983 |
| A3734 | Aracá | AR1 | 0.5556 | -54.4983 |
| A3746 | Aracá | AR2 | 0.5453 | -63.4539 |
| A3748 | Aracá | AR2 | 0.5453 | -63.4539 |
| A3750 | Aracá | AR2 | 0.5453 | -63.4539 |
| A3751 | Aracá | AR2 | 0.5453 | -63.4539 |
| A3752 | Aracá | AR2 | 0.5453 | -63.4539 |
| A3755 | Aracá | AR2 | 0.5453 | -63.4539 |
| A3759 | Aracá | AR2 | 0.5453 | -63.4539 |
| A3763 | Aracá | AR2 | 0.5453 | -63.4539 |
| A3764 | Aracá | AR2 | 0.5453 | -63.4539 |
| A3769 | Aracá | AR2 | 0.5453 | -63.4539 |
| A3772 | Aracá | AR2 | 0.5453 | -63.4539 |
| A3233 | Aracá | AR2 | 0.5453 | -63.4539 |
| A3777 | Aracá | AR3 | 0.6103 | -63.4303 |
| A3783 | Aracá | AR3 | 0.6103 | -63.4303 |
| A3784 | Aracá | AR3 | 0.6103 | -63.4303 |
| A3786 | Aracá | AR3 | 0.6103 | -63.4303 |
| A3787 | Aracá | AR3 | 0.6103 | -63.4303 |
| A3790 | Aracá | AR3 | 0.6103 | -63.4303 |
| A3791 | Aracá | AR3 | 0.6103 | -63.4303 |
| A3798 | Aracá | AR3 | 0.6103 | -63.4303 |
| A3799 | Aracá | AR3 | 0.6103 | -63.4303 |
| A3802 | Aracá | AR3 | 0.6103 | -63.4303 |
| A3805 | Aracá | AR3 | 0.6103 | -63.4303 |
| A3806 | Aracá | AR3 | 0.6103 | -63.4303 |
| A3234 | Aracá | AR3 | 0.6103 | -63.4303 |
| A3804 | Aracá | AR3 | 0.6103 | -63.4303 |
| A3830 | Aracá | AR4 | 0.4689 | -63.4756 |
| A3811 | Aracá | AR4 | 0.4689 | -63.4756 |
| A3835 | Aracá | AR4 | 0.4689 | -63.4756 |
| A3237 | Aracá | AR4 | 0.4689 | -63.4756 |
| A3240 | Aracá | AR4 | 0.4689 | -63.4756 |
| A3242 | Aracá | AR4 | 0.4689 | -63.4756 |
| A3248 | Aracá | AR4 | 0.4689 | -63.4756 |
| A3809 | Aracá | AR4 | 0.4689 | -63.4756 |
| A3810 | Aracá | AR4 | 0.4689 | -63.4756 |
| A3843 | Aracá | AR4 | 0.4689 | -63.4756 |
| A3860 | Aracá | AR5 | 0.4067 | -63.4092 |
| A3877 | Aracá | AR5 | 0.4067 | -63.4092 |
| A3881 | Aracá | AR5 | 0.4067 | -63.4092 |
| A3884 | Aracá | AR5 | 0.4067 | -63.4092 |
| A3889 | Aracá | AR5 | 0.4067 | -63.4092 |
| A3891 | Aracá | AR5 | 0.4067 | -63.4092 |
| A3894 | Aracá | AR5 | 0.4067 | -63.4092 |
| A3253 | Aracá | AR5 | 0.4067 | -63.4092 |
| A3872 | Aracá | AR5 | 0.4067 | -63.4092 |
| A3883 | Aracá | AR5 | 0.4067 | -63.4092 |
| A3898 | Aracá | AR5 | 0.4067 | -63.4092 |
| A3900 | Aracá | AR5 | 0.4067 | -63.4092 |
| A3922 | Aracá | AR7 | 0.3267 | -63.2622 |
| A3926 | Aracá | AR7 | 0.3267 | -63.2622 |
| A3929 | Aracá | AR7 | 0.3267 | -63.2622 |
| A3933 | Aracá | AR7 | 0.3267 | -63.2622 |
| A3943 | Aracá | AR7 | 0.3267 | -63.2622 |
| A3914 | Aracá | AR7 | 0.3267 | -63.2622 |
| A3915 | Aracá | AR7 | 0.3267 | -63.2622 |
| A3917 | Aracá | AR7 | 0.3267 | -63.2622 |
| A3923 | Aracá | AR7 | 0.3267 | -63.2622 |
| A3938 | Aracá | AR7 | 0.3267 | -63.2622 |
| A3841 | Aracá | AR7 | 0.3267 | -63.2622 |
| A3948 | Aracá | AR7 | 0.3267 | -63.2622 |
| A3949 | Aracá | AR7 | 0.3267 | -63.2622 |
| A3952 | Aracá | AR7 | 0.3267 | -63.2622 |
| A4295 | Uatumã | UT1 | -2.2817 | -59.0617 |
| A4296 | Uatumã | UT1 | -2.2817 | -59.0617 |
| A4298 | Uatumã | UT1 | -2.2817 | -59.0617 |
| A4299 | Uatumã | UT1 | -2.2817 | -59.0617 |
| A4301 | Uatumã | UT1 | -2.2817 | -59.0617 |
| A4303 | Uatumã | UT1 | -2.2817 | -59.0617 |
| A4310 | Uatumã | UT1 | -2.2817 | -59.0617 |
| A4335 | Uatumã | UT1 | -2.2817 | -59.0617 |
| A4343 | Uatumã | UT1 | -2.2817 | -59.0617 |
| A4336 | Uatumã | UT1 | -2.2817 | -59.0617 |
| A3992 | Uatumã | UT10 | -2.2858 | -58.865 |
| A3993 | Uatumã | UT10 | -2.2858 | -58.865 |
| A3994 | Uatumã | UT10 | -2.2858 | -58.865 |
| A3995 | Uatumã | UT10 | -2.2858 | -58.865 |
| A3998 | Uatumã | UT10 | -2.2858 | -58.865 |
| A3999 | Uatumã | UT10 | -2.2858 | -58.865 |
| A4006 | Uatumã | UT10 | -2.2858 | -58.865 |
| A4009 | Uatumã | UT10 | -2.2858 | -58.865 |
| A4011 | Uatumã | UT10 | -2.2858 | -58.865 |
| A3961 | Uatumã | UT12 | -2.2728 | -58.6719 |
| A3975 | Uatumã | UT12 | -2.2728 | -58.6719 |
| A3877 | Uatumã | UT12 | -2.2728 | -58.6719 |
| A3981 | Uatumã | UT12 | -2.2728 | -58.6719 |
| A3273 | Uatumã | UT12 | -2.2728 | -58.6719 |
| A3**2**87 | Uatumã | UT12 | -2.2728 | -58.6719 |
| A3291 | Uatumã | UT12 | -2.2728 | -58.6719 |
| A3298 | Uatumã | UT12 | -2.2728 | -58.6719 |
| A3971 | Uatumã | UT12 | -2.2728 | -58.6719 |
| A4279 | Uatumã | UT2 | -2.2867 | -58.9561 |
| A4280 | Uatumã | UT2 | -2.2867 | -58.9561 |
| A4282 | Uatumã | UT2 | -2.2867 | -58.9561 |
| A4338 | Uatumã | UT2 | -2.2867 | -58.9561 |
| A4339 | Uatumã | UT2 | -2.2867 | -58.9561 |
| A4340 | Uatumã | UT2 | -2.2867 | -58.9561 |
| A4342 | Uatumã | UT2 | -2.2867 | -58.9561 |
| A4361 | Uatumã | UT2 | -2.2867 | -58.9561 |
| A4362 | Uatumã | UT2 | -2.2867 | -58.9561 |
| A4363 | Uatumã | UT2 | -2.2867 | -58.9561 |
| A4281 | Uatumã | UT2 | -2.2867 | -58.9561 |
| A4349 | Uatumã | UT2 | -2.2867 | -58.9561 |
| A4402 | Uatumã | UT3 | -2.2822 | -59.03 |
| A4403 | Uatumã | UT3 | -2.2822 | -59.03 |
| A4405 | Uatumã | UT3 | -2.2822 | -59.03 |
| A4406 | Uatumã | UT3 | -2.2822 | -59.03 |
| A4415 | Uatumã | UT3 | -2.2822 | -59.03 |
| A4417 | Uatumã | UT3 | -2.2822 | -59.03 |
| A4424 | Uatumã | UT3 | -2.2822 | -59.03 |
| A4436 | Uatumã | UT3 | -2.2822 | -59.03 |
| A4443 | Uatumã | UT3 | -2.2822 | -59.03 |
| A4445 | Uatumã | UT3 | -2.2822 | -59.03 |
| A4407 | Uatumã | UT3 | -2.2822 | -59.03 |
| A4422 | Uatumã | UT3 | -2.2822 | -59.03 |
| A4439 | Uatumã | UT3 | -2.2822 | -59.03 |
| A4368 | Uatumã | UT5 | -2.1817 | -59.0217 |
| A4373 | Uatumã | UT5 | -2.1817 | -59.0217 |
| A4374 | Uatumã | UT5 | -2.1817 | -59.0217 |
| A4383 | Uatumã | UT5 | -2.1817 | -59.0217 |
| A4388 | Uatumã | UT5 | -2.1817 | -59.0217 |
| A6625 | Viruá | V1 | 1.4093 | -60.9912 |
| A6630 | Viruá | V1 | 1.4093 | -60.9912 |
| A6639 | Viruá | V1 | 1.4093 | -60.9912 |
| A6647 | Viruá | V1 | 1.4093 | -60.9912 |
| A6650 | Viruá | V1 | 1.4093 | -60.9912 |
| A6651 | Viruá | V1 | 1.4093 | -60.9912 |
| A6681 | Viruá | V1 | 1.4093 | -60.9912 |
| A6689 | Viruá | V1 | 1.4093 | -60.9912 |
| A6690 | Viruá | V1 | 1.4093 | -60.9912 |
| A6692 | Viruá | V1 | 1.4093 | -60.9912 |
| A6693 | Viruá | V1 | 1.4093 | -60.9912 |
| A6708 | Viruá | V1 | 1.4093 | -60.9912 |
| A6805 | Viruá | V3 | 1.4317 | -60.8684 |
| A6807 | Viruá | V3 | 1.4317 | -60.8684 |
| A6810 | Viruá | V3 | 1.4317 | -60.8684 |
| A6811 | Viruá | V3 | 1.4317 | -60.8684 |
| A6814 | Viruá | V3 | 1.4317 | -60.8684 |
| A6816 | Viruá | V3 | 1.4317 | -60.8684 |
| A6861 | Viruá | V3 | 1.4317 | -60.8684 |
| A6883 | Viruá | V3 | 1.4317 | -60.8684 |
| A6894 | Viruá | V4 | 1.5962 | -61.0456 |
| A6909 | Viruá | V4 | 1.5962 | -61.0456 |
| A6912 | Viruá | V4 | 1.5962 | -61.0456 |
| A10204 | Viruá | V4 | 1.5962 | -61.0456 |
| A10205 | Viruá | V4 | 1.5962 | -61.0456 |
| A10206 | Viruá | V4 | 1.5962 | -61.0456 |
| A10209 | Viruá | V4 | 1.5962 | -61.0456 |
| A10210 | Viruá | V4 | 1.5962 | -61.0456 |
| A10711 | Viruá | V4 | 1.5962 | -61.0456 |
| A10714 | Viruá | V4 | 1.5962 | -61.0456 |
| A6758 | Viruá | V5 | 1.6621 | -60.9325 |
| A6759 | Viruá | V5 | 1.6621 | -60.9325 |
| A6769 | Viruá | V5 | 1.6621 | -60.9325 |
| A6773 | Viruá | V5 | 1.6621 | -60.9325 |
| A6786 | Viruá | V5 | 1.6621 | -60.9325 |
| A6787 | Viruá | V5 | 1.6621 | -60.9325 |
| A6788 | Viruá | V5 | 1.6621 | -60.9325 |
| A6789 | Viruá | V5 | 1.6621 | -60.9325 |
| A6790 | Viruá | V5 | 1.6621 | -60.9325 |
| A6791 | Viruá | V5 | 1.6621 | -60.9325 |
| A10153 | Viruá | V5 | 1.6621 | -60.9325 |
| A10172 | Viruá | V5 | 1.6621 | -60.9325 |
| A10176 | Viruá | V5 | 1.6621 | -60.9325 |
| A10177 | Viruá | V5 | 1.6621 | -60.9325 |
| A10179 | Viruá | V5 | 1.6621 | -60.9325 |
| A10188 | Viruá | V5 | 1.6621 | -60.9325 |
| A10189 | Viruá | V5 | 1.6621 | -60.9325 |
| A10197 | Viruá | V5 | 1.6621 | -60.9325 |

**Table S2** - Number of alleles (N_A_), allelic richness (A_R_), observed (H_o_) and expected (H_e_) heterozygosity, and Wright’s fixation index (F_IS_) of 15 microsatellite loci within *E. ruficeps* populations.

|  | **Aracá** | | | | | **Uatumã** | | | | | **Viruá** | | | | |
| --- | --- | --- | --- | --- | --- | --- | --- | --- | --- | --- | --- | --- | --- | --- | --- |
|  | **N_A_** | **A_R_** | **H_o_** | **H_e_** | **F_IS_** | **N_A_** | **A_R_** | **H_o_** | **H_e_** | **F_IS_** | **N_A_** | **A_R_** | **H_o_** | **H_e_** | **F_IS_** |
| **Loc1** | 14 | 14.774 | 51 | 61.175 | 0.167 | 14 | 14.559 | 37.000 | 48.982 | 0.246 | 14 | 14.000 | 35.000 | 43.116 | 0.190 |
| **Loc2** | 4 | 3.887 | 33.000 | 26.958 | -0.226 | 5 | 4.939 | 26.000 | 27.800 | 0.065 | 5 | 5.000 | 19.000 | 18.526 | -0.026 |
| **Loc3** | 6 | 6.222 | 17.000 | 18.190 | 0.066 | 5 | 4.799 | 23.000 | 26.435 | 0.131 | 3 | 3.000 | 10.000 | 17.558 | 0.4331 |
| **Loc4** | 10 | 9.556 | 53.000 | 54.631 | 0.030 | 9 | 9.445 | 33.000 | 40.451 | 0.186 | 6 | 6.000 | 28.000 | 33.295 | 0.160 |
| **Loc5** | 19 | 18.727 | 33.000 | 59.225 | 0.445 | 26 | 25.726 | 39.000 | 44.215 | 0.1191 | 18 | 19.000 | 24.000 | 38.161 | 0.374 |
| **Loc6** | 5 | 4.333 | 32.000 | 36.000 | 0.1118 | 5 | 4.943 | 32.000 | 32.896 | 0.0275 | 5 | 5.000 | 24.000 | 24.947 | -0.124 |
| **Loc7** | 3 | 3.557 | 17.000 | 16.702 | -0.0180 | 2 | 2.972 | 11.000 | 10.200 | 0.028 | 3 | 4.000 | 19.000 | 19.226 | 0.012 |
| **Loc8** | 15 | 13.550 | 47.000 | 47.237 | 0.005 | 12 | 11.909 | 45.000 | 37.047 | -0.217 | 8 | 9.000 | 32.000 | 30.312 | -0.0563 |
| **Loc9** | 5 | 5.620 | 33.000 | 30.259 | -0.091 | 8 | 8.277 | 12.000 | 14.505 | 0.174 | 5 | 5.000 | 23.000 | 23.200 | 0.009 |
| **Loc10** | 14 | 13.545 | 44.000 | 51.714 | 0.150 | 9 | 9.798 | 30.000 | 36.047 | 0.169 | 12 | 12.000 | 20.000 | 34.325 | 0.4205 |
| **Loc11** | 6 | 5.318 | 21.000 | 21.413 | 0.0194 | 7 | 6.282 | 19.000 | 18.939 | -0.003 | 4 | 4.000 | 25.000 | 23.337 | -0.0721 |
| **Loc12** | 28 | 25.841 | 64.000 | 64.148 | 0.0023 | 30 | 28.917 | 51.000 | 53.735 | 0.051 | 21 | 22.000 | 26.000 | 35.120 | 0.262 |
| **Loc13** | 7 | 7.114 | 18.000 | 35.899 | 0.501 | 5 | 5.971 | 14.000 | 26.545 | 0.475 | 9 | 10.000 | 14.000 | 29.966 | 0.536 |
| **Loc14** | 21 | 20.067 | 66.000 | 63.820 | -0.034 | 19 | 19.578 | 49.000 | 51.982 | 0.0579 | 19 | 20.000 | 42.000 | 42.253 | 0.006 |
| **Loc15** | 4 | 4.890 | 10.000 | 32.840 | 0.697 | 6 | 6.770 | 9.000 | 19.371 | 0.538 | 5 | 6.000 | 9.000 | 18.217 | 0.7284 |

**Table S3** – Resistance values by landscape category attributed by each expert Amazon Ornithologist and the average used in the resistance models.

| **Land Cover** | **Expert1** | **Expert2** | **Expert3** | **Expert4** | **Average** |
| --- | --- | --- | --- | --- | --- |
| Campina (open white-sand vegetation) | 0.01 | 0.01 | 0.1 | 0.01 | 0.04 |
| Campinarana (forests white-sand vegetation) | 0.60 | 0.1 | 0.3 | 0.1 | 0.16667 |
| Igapó (black-water flooded forest) | 0.60 | 0.15 | 0.4 | 0.25 | 0.26667 |
| Várzea (white-water flooded forest) | 0.99 | ?? | 0.4 | 0.8 | 0.6 |
| Terra firme forest | 0.99 | 0.8 | 0.7 | 0.99 | 0.83 |
| Deforested área (anthropogenic) | 0.30 | 0.8 | 0.5 | 0.75 | 0.68333 |
| Big black-water river  (e.g. Negro River) | 0.6 | 0.5 | 0.6 | 0.5 | 0.53333 |
| Big white-water river (e.g. Amazon River) | 0.99 | ?? | 0.7 | 0.99 | 0.845 |
| Medium black-water river  (e.g. AracáRiver) | 0.10 | 0.25 | 0.8 | 0.25 | 0.43333 |
| Medium white-water river (e.g. Branco River) | 0.60 | ?? | 0.8 | 0.8 | 0.8 |

**Table S4** - The genetic diversity measurements for each individual landscape. Nucleotide (Pi) and haplotype (H_D_) diversity from mitochondrial data, and allelic richness (N_G_) and genetic diversity (Theta) from microsatellite data.

|  | **ND2** | | **Microsat** | |
| --- | --- | --- | --- | --- |
| **Landscape** | **P_i_** | **H_D_** | **N_G_** | **Theta** |
| **Aracá** | **0.003 +/- 0.0006** | **0.84 +/- 0.07** | **24 +/- 3.58** | 1.61 +/-0.06 |
| **Uatumã** | 0.001 +/- 0.0004 | 0.78 +/- 0.1 | 19.33 +/- 5.61 | 1.56 +/- 0.05 |
| **Viruá** | 0.001 +/- 0.0003 | 0.76 +/- 0.04 | 24 +/- 8.64 | **1.69 +/- 0.04** |

**Table S5** – F_ST_ from ND2 sequences data between the sites inside and among each landscape. AR represents sites in Aracá, UT in Uatumã and V in Viruá.

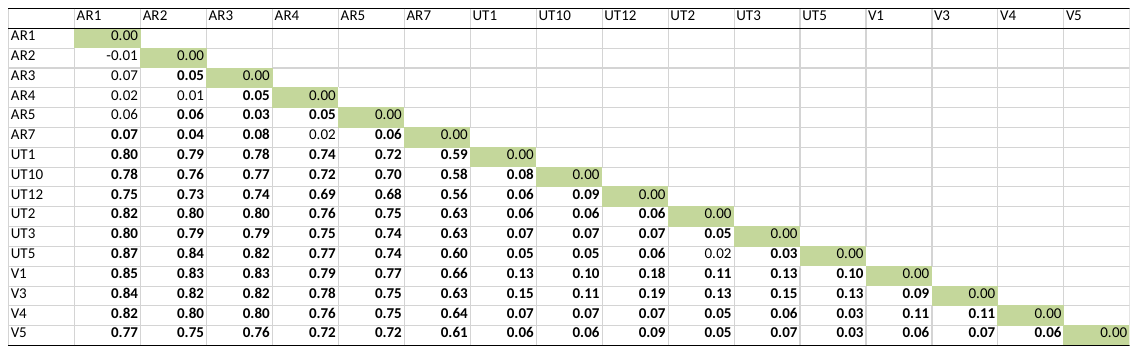

**Table S6** – F_ST_ from microsatellite data between the sites inside and among landscape. Significant values are in bold. AR represents sites in Aracá, UT in Uatumã and V in Viruá.

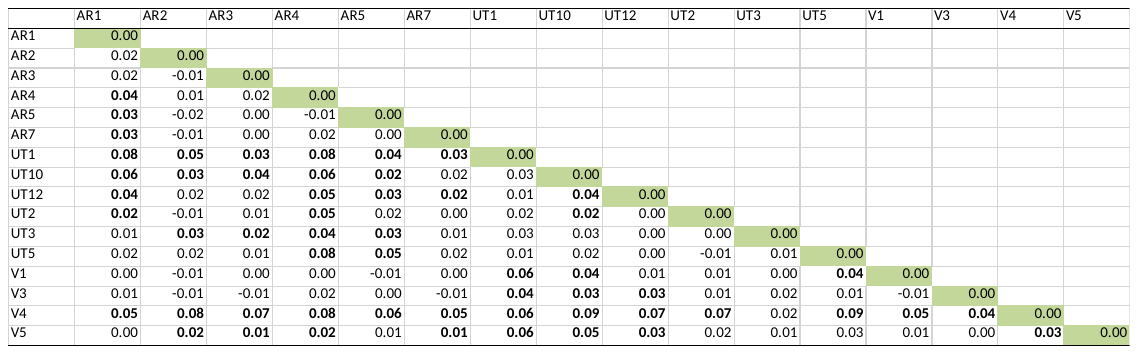

**Table S7** – Estimated parameters (values estimated with standardize error, t-value and respective p-value) of the best fit models selected in model selection. The genetic diversity variables for mitochondrial data are nucleotide (Pi) and haplotype (H_D_) diversity and for the microsatellite data are Theta and N_G_. For Pi and Theta just landscape was selected to explain diversity. HD the best model was the constant null model. For N_G_ the best model was just with the proximity index.

| **Variable** | **Parameters** | **Estimate** | **Std. Error** | **t value** | **Pr(>\|t\|)** |
| --- | --- | --- | --- | --- | --- |
| **Pi** | **Intercept** | 0.0032333 | 0.0001941 | 16.654 | 3.77E-10 |
|  | **Uatuma** | -0.0018567 | 0.0002746 | -6.762 | 1.34E-05 |
|  | **Virua** | -0.0019808 | 0.000307 | -6.453 | 2.16E-05 |
| **H_D_** | **Intercept** | 0.7945 | 0.02029 | 39.16 | <2e-16 |
| **Theta** | **Intercept** | 1.60514 | 0.02082 | 77.103 | <2e-16 |
|  | **Uatuma** | -0.04426 | 0.02944 | -1.503 | 0.1567 |
|  | **Virua** | 0.08194 | 0.03292 | 2.489 | 0.0271 |
| **N_G_** | **Intercept** | 1.97E+01 | 1.35E+00 | 14.604 | 7.25E-10 |
|  | **Proximity index** | 1.50E-03 | 4.31E-04 | 3.487 | 0.00363 |

**Table S8** – Mantel results for the sites inside each landscape for test of isolation by distance and isolation by resistance. The response variables are the F_ST_.

| **Isolation** | **Landscape** | **Micros** | | **ND2** | |
| --- | --- | --- | --- | --- | --- |
|  |  | **R** | **p** | **R** | **P** |
| **Distance** | **Aracá** | 0.26 | 0.3 | -0.41 | 0.93 |
|  | **Viruá** | 0.04 | 0.42 | -0.2 | 0.83 |
|  | **Uatumã** | 0.002 | 0.57 | 0.13 | 0.24 |
| **Resistence** | **Aracá** | 0.21 | 0.31 | 0.005 | 0.5 |
|  | **Viruá** | -0.14 | 0.7 | -0.56 | 0.92 |
|  | **Uatumã** | 0.11 | 0.4 | 0.33 | 0.19 |

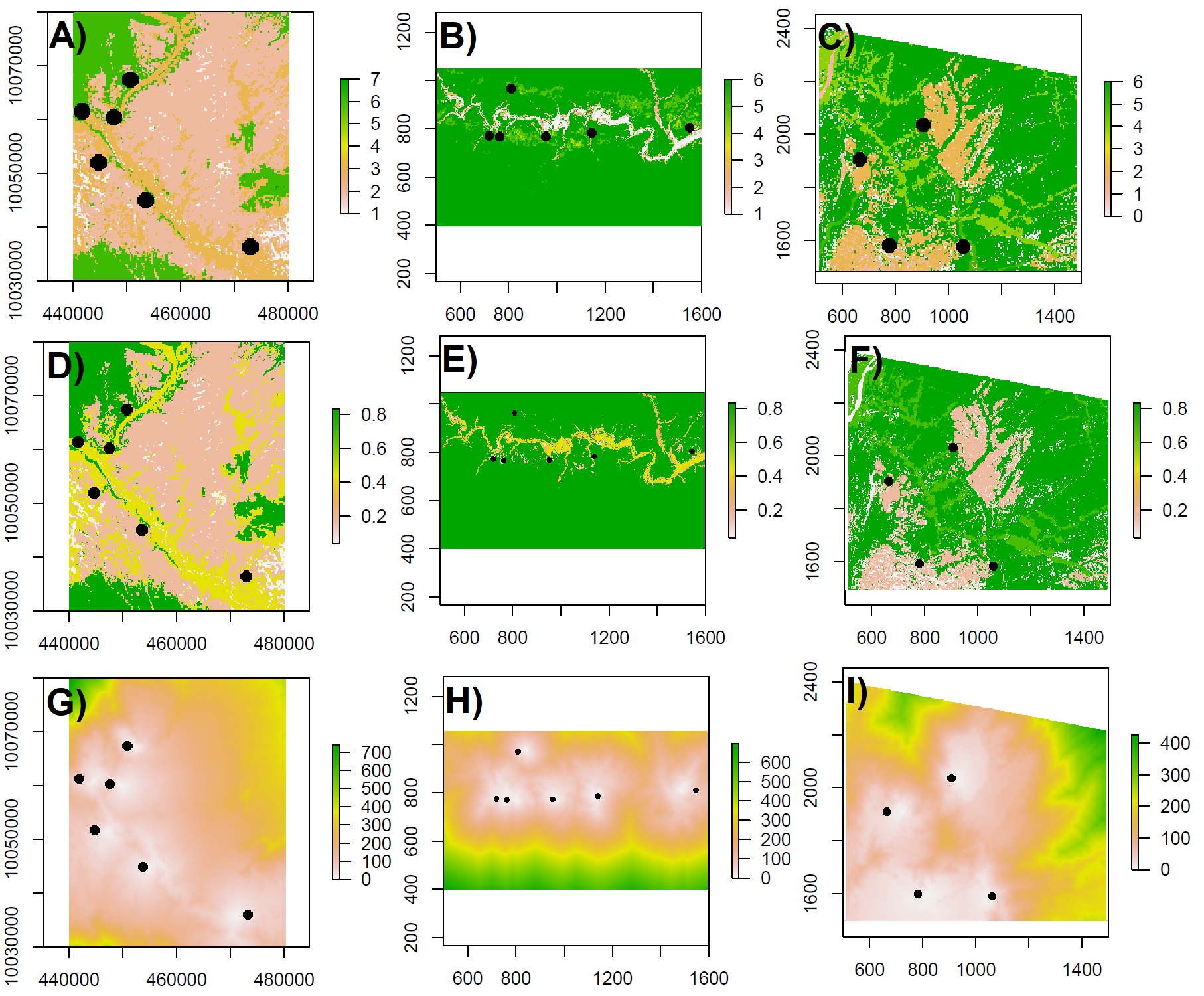

**Figure S1.** Raster layers by landscape used to create the resistance matrix. Classified raster for vegetation category from A) Aracá, B) Uatumã and, C) Viruá. The classified raster by resistance matrix from D) Aracá, E) Uatumã and, F) Viruá. The conductance transitional layer created to measure the resistance between the pairwise points from G) Aracá, H) Uatumã and, I) Viruá.

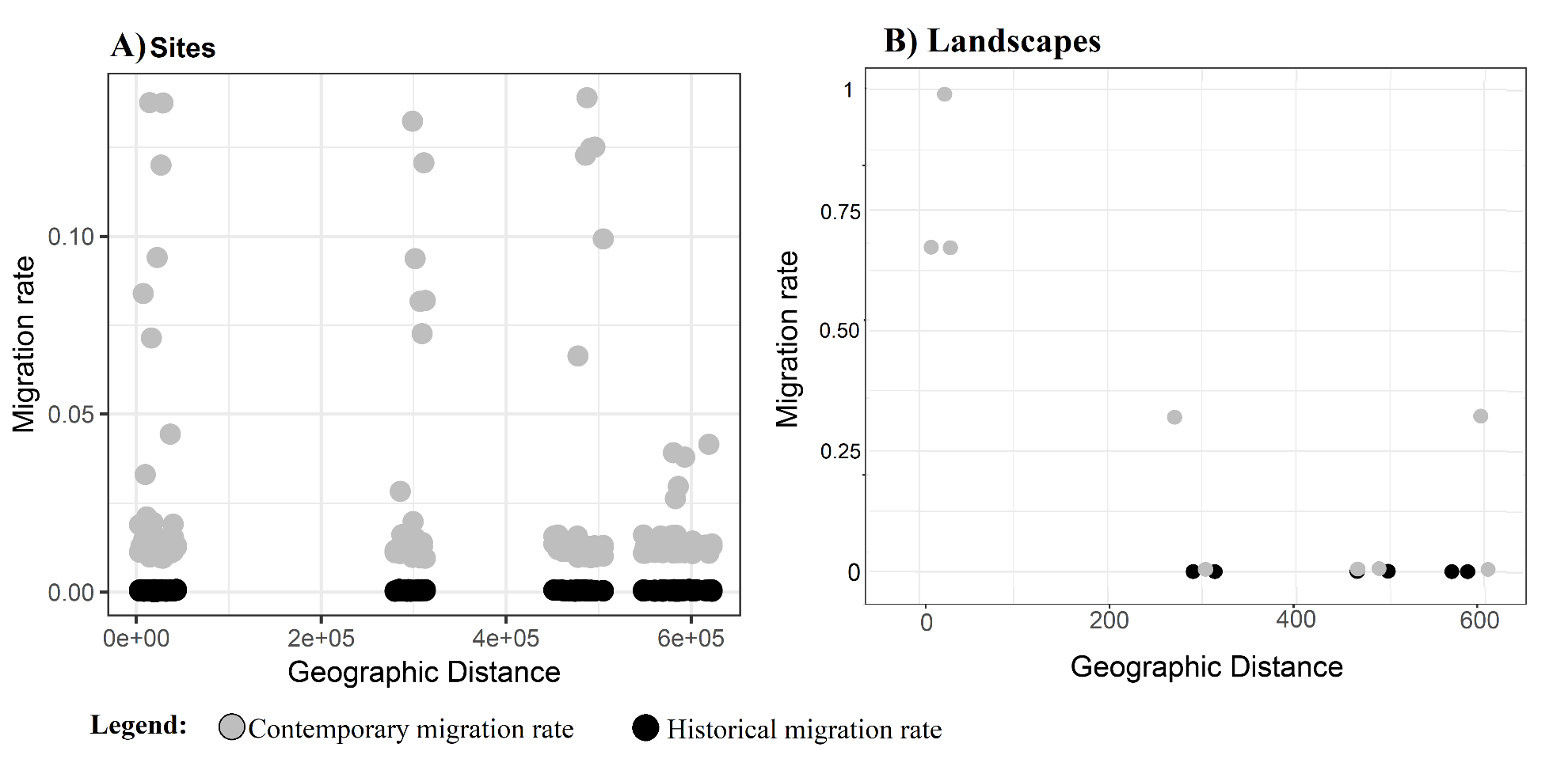

**Figure S2.** Spatial correlation between migration rates and geographic distances in A) all sites, and B) among landscape. Gray points represent the contemporary migration rate calculate from microsatellite data using the BayesAss software. Black points and line represent the historical migration rate calculated from ND2 sequences, both estimated by Migrate-N software.

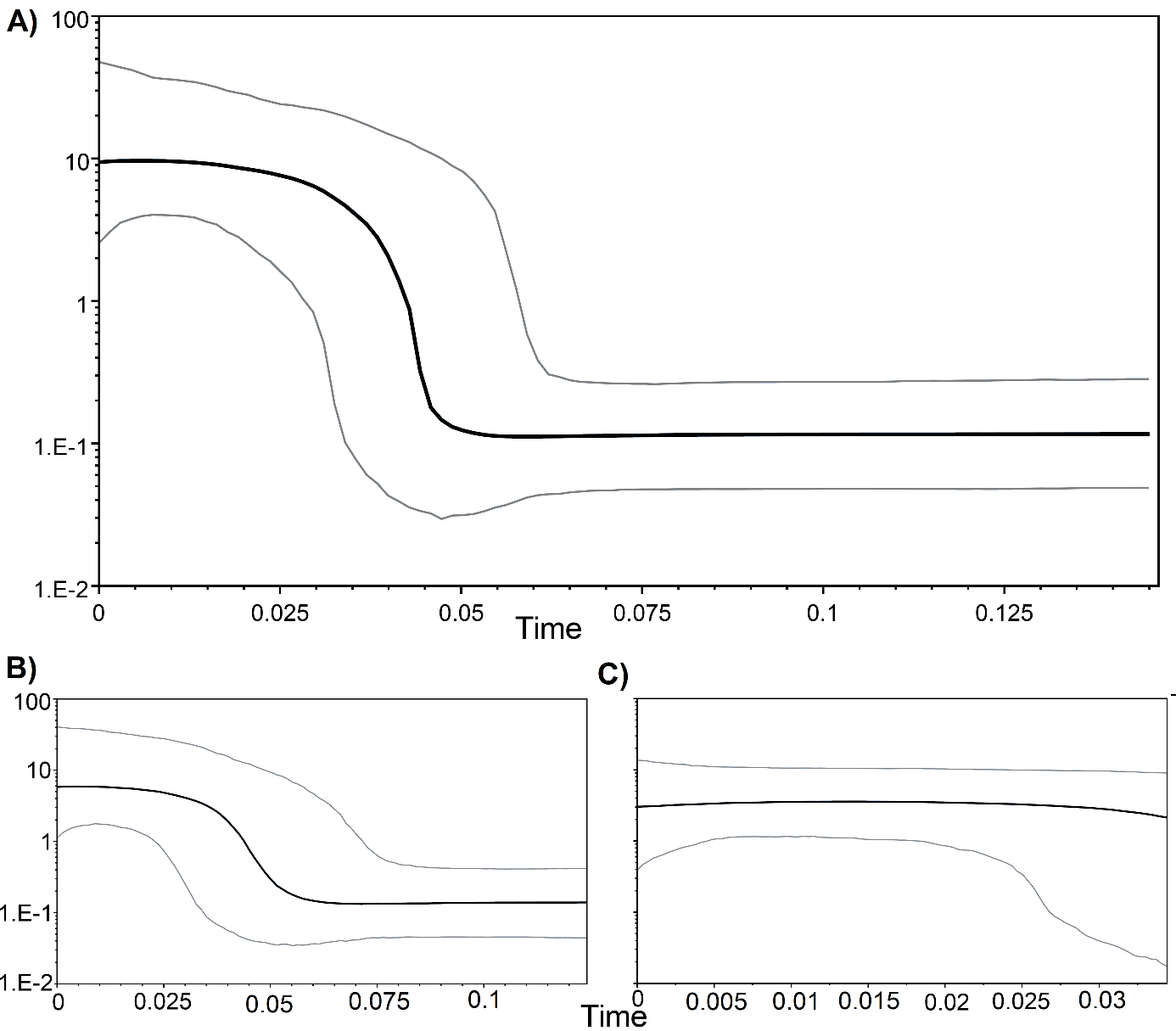

**Figure S3**. Bayesian skyline plot based on 978 bp of 178 mitochondrial ND2 sequences showing the demographic history of sampled *Elaenia ruficeps* populations in the study region (A). (B) Is the population demography from Aracá, and (C) from Viruá + Uatumã. The horizontal axis shows time in thousands of years before the present and the vertical axis shows effective population size. The black line represents the median value, while the gray lines indicate the 95% Bayesian credibility intervals.
